# Task-based attentional and default mode connectivity associated with STEM anxiety profiles among university physics students

**DOI:** 10.1101/2022.09.30.508557

**Authors:** Donisha D. Smith, Alan Meca, Katherine L. Bottenhorn, Jessica E. Bartley, Michael C. Riedel, Taylor Salo, Julio A. Peraza, Robert W. Laird, Shannon M. Pruden, Matthew T. Sutherland, Eric Brewe, Angela R. Laird

**Author notes:** Corresponding Author*: **Dr. Angela R. Laird**, Professor of Physics, Florida International University, Miami, FL, USA |.

## Abstract

Attentional control theory (ACT) posits that elevated anxiety increases the probability of re-allocating cognitive resources needed to complete a task to processing anxiety-related stimuli. This process impairs processing efficiency and can lead to reduced performance effectiveness. Science, technology, engineering, and math (STEM) students frequently experience STEM-related anxiety, which can interfere with learning and performance and negatively impact student retention and graduation rates. The objective of this study was to extend the ACT framework to investigate the neurobiological associations between STEM-related anxiety and cognitive performance among 123 physics undergraduate students. Latent profile analysis (LPA) identified four profiles of student STEM-related anxiety, including two profiles that represented the majority of the sample (Low STEM Anxiety; 59.3% and High Math Anxiety; 21.9%) and two additional profiles that were not well represented (High STEM Anxiety; 6.5% and High Science Anxiety; 4.1%). Students underwent a functional magnetic resonance imaging (fMRI) session in which they performed two tasks involving physics cognition: the Force Concept Inventory (FCI) task and the Physics Knowledge (PK) task. No significant differences were observed in FCI or PK task performance between High Math Anxiety and Low STEM Anxiety students. During the three phases of the FCI task, we found no significant brain connectivity differences during scenario and question presentation, yet we observed significant differences during answer selection within and between the dorsal attention network (DAN), ventral attention network (VAN), and default mode network (DMN). Further, we found significant group differences during the PK task were limited to the DAN, including DAN-VAN and within-DAN connectivity. These results highlight the different cognitive processes required for physics conceptual reasoning compared to physics knowledge retrieval, provide new insight into the underlying brain dynamics associated with anxiety and physics cognition, and confirm the relevance of ACT theory for STEM-related anxiety.

## Introduction

Improving student retention rates among science, technology, engineering, and math (STEM) university majors has been an enduring issue in higher education (Almatrafi et al., 2017; Findley-Van Nostrand & Pollenz, 2017). Only ~40% of university undergraduate students enrolled in STEM degree programs in the United States complete their degree (Waldrop, 2015), yet within the next 10-20 years a projected one million STEM-related jobs will need to be filled by qualified individuals (Findley-Van Nostrand & Pollenz, 2017). These low STEM retention rates among U.S. students have prompted multiple research studies and programmatic initiatives dedicated to investigating and addressing the motivational, institutional, and cognitive factors that result in students abandoning STEM degree programs (Cromley et al., 2016; Leary et al., 2020). While findings suggest that STEM retention is a multifaceted problem, one notable psychosocial barrier that students commonly report facing when choosing whether or not to remain in their programs is STEM-related anxiety, which is defined as apprehension or fear towards STEM-related activities (Suárez-Pellicioni et al., 2016). STEM-related anxiety has been associated with underperformance in STEM courses (Daker et al., 2021; Sithole et al., 2017), avoidance of effortful and effective study strategies (Jenifer et al., 2022), and is a significant contributing factor to withdrawal from introductory university STEM courses (Daker et al., 2021).

The association between STEM-related anxiety and diminished STEM performance may be explained by Attentional Control Theory (ACT), which posits that elevated anxiety impairs efficient functioning by reducing cognitive resources available for attentional focus, thereby compromising performance effectiveness (Corbetta & Shulman, 2002; Derakshan et al., 2009; Eysenck et al., 2007; Suárez-Pellicioni et al., 2016). ACT is an updated adaptation of the processing efficiency theory proposed by Eysenck & Calvo (1992). Processing efficiency theory distinguishes between performance effectiveness, which is defined as the quality of performance on a task, and processing efficiency, which moderates the relation between performance efficiency and the cognitive resources needed to attain a particular quality of performance. Processing efficiency theory also proposes that excessive rumination among highly anxious individuals encourages them to exert more cognitive effort to compensate for the deleterious effects of anxiety. Elevated use of finite cognitive resources is thought to lead to highly anxious individuals displaying reduced processing efficiency, which in turn worsens performance efficiency. Eysenck et al. (2005) provided experimental evidence supporting this theory by demonstrating that groups of individuals with high and low anxiety performed similarly when a primary, visuospatial short-term memory task was coupled with a secondary, simple motor tapping task. However, when the secondary task was altered to require the use of the working memory system (e.g., counting backwards), highly anxious individuals performed worse on both the primary and secondary tasks, thus suggesting that elevated anxiety levels may diminish processing efficiency, thereby degrading performance efficiency (Eysenck et al., 2005).

While processing efficiency theory broadly predicts that anxiety impairs executive functioning, which includes multiple cognitive processes ranging from attention shifting to updating working memory (Derakshan & Eysenck, 2009), ACT narrows the scope to posit that anxiety specifically impairs attentional control (Corbetta & Shulman, 2002). Within this framework, increased anxiety is thought to disrupt the equilibrium between two attentional systems: a goal-directed dorsal frontoparietal system, often referred to as the dorsal attention network (DAN) (Corbetta et al., 2008; Corbetta & Shulman, 2002; Fox et al., 2006; Hacker et al., 2017), which is engaged during task-relevant processes, and a stimulus-driven ventral parietal system, often referred to as the ventral attention network (VAN) (Corbetta et al., 2008; Corbetta & Shulman, 2002; Fox et al., 2006; Hacker et al., 2017), which is involved in the processing of rewarding and aversive stimuli (Derakshan & Eysenck, 2009; Dosenbach et al., 2007; Lydon-Staley et al., 2019; Vossel et al., 2014). According to ACT, the detrimental effects of increased anxiety can be alleviated by increased activation of stimulus-driven attentional systems. However, as a consequence of the elevated activity of stimulus-driven attentional systems, fewer cognitive resources are dedicated to goal-directed attentional systems. In anxious individuals, enhanced activation of stimulus-driven attentional systems and decreases in goal-directed attentional systems are thought to occur in tandem, reflecting an aversive, elevated tendency to flight responses which lead to reduced performance and processing ability (Corbetta & Shulman, 2002). This mechanism may result in a feedback loop where elevated anxiety and reduced performance lead to lowered self-efficacy, leading to maintenance of disequilibrium between goal-directed and stimulus-driven attentional systems (Suárez-Pellicioni et al., 2016). Furthermore, while ACT emphasizes both the independent and dependent functionalities of the DAN and VAN, evidence suggests that both engagement and disengagement of the default mode network (DMN) also influences attentional control and affects task performance (Poole et al., 2016). The DMN is often categorized as the ‘task-negative’ network since it is often suppressed during cognitively demanding tasks and active during periods of self-referential processing and mind-wandering (i.e., in the absence of tasks) (Alves et al., 2019; Fortenbaugh et al., 2017). Prior work has demonstrated that interdependence within and between regions of the DMN, DAN, and VAN are associated with variability in task performance (Anticevic et al., 2012; Elton & Gao, 2015; Kelly et al., 2008), particularly during sustained attention tasks (Fortenbaugh et al., 2017; Kucyi et al., 2016). Furthermore, DMN activity alterations among anxious individuals (Qiao et al., 2020) suggests that both within-network activation of the DMN and its interactions with both the DAN and VAN are important to attentional control and relevant to ACT (Poole et al., 2016).

The objective of the present study was to extend the ACT framework to study the neurobiological associations between STEM-related anxiety and cognitive performance. Although the ACT framework has primarily been applied to the study of generalized anxiety, a recent meta-analysis demonstrated that math anxiety negatively impacts attentional control (Finell et al., 2022), confirming the relevance of ACT for STEM-related anxiety. Towards this end, we investigated STEM anxiety among undergraduate students enrolled in an introductory physics course. Students participated in a behavioral session in which they completed self-reports of STEM (i.e., science and math) anxiety, followed by a functional magnetic resonance imaging (fMRI) session in which they performed two tasks involving physics cognition: the Force Concept Inventory task (FCI) and the Physics Knowledge (PK) task. Latent profile analysis (LPA) was used to identify groups of students with similar STEM anxiety profiles. Measures of between- and within-network connectivity were extracted from the DAN, VAN, and DMN networks during the FCI and PK tasks. Regression analyses were then conducted to determine task-based connectivity differences between STEM anxiety groups. We hypothesized that significant DAN- and VAN-related connectivity differences across STEM anxiety groups would be observed for both the FCI and PK tasks. However, since the FCI task involves sustained physics cognition, we further hypothesized that DMN-related differences would be observed only for the FCI task and not the PK task. Lastly, we expected to observe significant differences in task performance (i.e., accuracy) across STEM anxiety groups, as predicted by ACT and processing efficiency theory. Together, these findings may inform knowledge on the behavioral and neurobiological associations between STEM-related anxiety and physics cognition among undergraduate STEM students.

## Methods

### Participants

The study sample included 123 healthy, right-handed undergraduate students (mean age = 19.8 ± 1.5, range = 18-26 years; 56 females). Students were enrolled in introductory, calculus-based physics courses at Florida International University (FIU) in Miami, Florida. At enrollment, participants provided demographic information, such as their age, sex, ethnicity (i.e., Hispanic or non-Hispanic), household income, grade point average (GPA), and number of years enrolled as a student at FIU (i.e., freshman, sophomore, junior, or senior) (**Table 1**). Participants self-reported that they were free from cognitive impairments, neurological and psychiatric conditions, and did not use psychotropic medications.

**Table 1.** Participant Demographic Information.

|  | N | Percentage |
| --- | --- | --- |
| **Gender** |  |  |
| Male | 67 | 54 |
| Female | 56 | 46 |
| **Ethnicity** |  |  |
| Hispanic | 85 | 69 |
| Non-Hispanic | 38 | 31 |
| **Household Income** |  |  |
| < \$15,000 | 26 | 21 |
| \$15,000 - \$34,999 | 24 | 20 |
| \$35,000 - \$49,999 | 16 | 13 |
| \$50,000 - \$74,999 | 21 | 17 |
| \$75,000 - \$99,999 | 18 | 15 |
| >\$100,000 | 18 | 15 |
| **Years Enrolled** |  |  |
| Freshman | 11 | 10 |
| Sophomore | 51 | 47 |
| Junior | 33 | 31 |
| Senior | 13 | 12 |
|  | **Mean (Std. Dev.)** | **Range** |
| **Age** | 19.8 (1.5) | 18-26 |
| **GPA** | 3.3 (0.5) | 0.0-4.0 |

*Note.* The “N” column represents the sample size of the group and the “Percentage” column represents the percentage of participants in that group for each categorical variable (Gender, Ethnicity, Household Income, and Years Enrolled).

### Procedures

At the beginning of the semester, participant recruitment began with research assistants visiting eligible classrooms and delivering a brief presentation, with the permission of the professor, informing students about the opportunity to voluntarily participate in this study. Enrolled participants completed a behavioral and fMRI session at the beginning of the course (i.e., pre-instruction), no later than the fourth week of instruction and prior to the first course exam. Behavioral sessions were conducted in an on-campus lab and students were asked to complete a battery of Qualtrics surveys. Imaging sessions were conducted off-campus and participants were provided with free parking and/or FIU-organized transportation to and from the MRI site. Written informed consent was obtained in accordance with FIU’s Institutional Review Board approval. Participants were compensated monetarily after both the behavioral and fMRI sessions.

### STEM-Related Anxiety Measures

Participants completed a series of self-report instruments during their behavioral sessions, including, but not limited to, assessments of their STEM-related anxiety. The Science Anxiety Questionnaire (Mallow, 1994) consists of 22 items and had a Cronbach’s alpha of *α* = 0.86, which was calculated using the `cronbach.alpha` command from the `ltm` package available in R. The items asked students to indicate their level of discomfort with respect to a range of science-related subjects and activities (e.g., *“Having your professor watch you perform an experiment in the lab.”*) on a 5-point Likert scale, with “0” suggesting no apprehension and “4” indicating the highest level of discomfort. The Mathematics Anxiety Rating Scale (Alexander & Martray, 1989) consists of 25 items and had a Cronbach’s alpha of *α* = 0.96. The items asked students to indicate their level of discomfort with respect to a variety of mathematics related activities (e.g., *“Being given a set of division problems to solve on paper.”*) on a 5-point Likert scale, with “0” suggesting no apprehension and “4” indicating the highest level of discomfort.

### Generalized Anxiety Measure

In addition to these two measures of STEM-related anxiety, symptoms of anxiety were assessed to capture students’ self-report of non-STEM-related anxiety. The Beck Anxiety Inventory (Beck et al., 1988) is a Likert scale consisting of 21 items and had a Cronbach’s alpha of *α* = 0.94. The items asked students to indicate the severity of anxiety related symptoms that they experienced within the past month (e.g., “*Fear of the worst happening*”) on a 4-point Likert scale with “0” indicating that a symptom hadn’t been experienced in the past month and “3” indicating that the symptom had been severe in the past month.

### MRI Data Acquisition

MRI data were acquired on a GE 3T Healthcare Discovery 750W MRI scanner at the University of Miami. Functional imaging data were acquired with an interleaved gradient-echo, echo planar imaging (EPI) sequence (TR/TE = 2000/30ms, flip angle = 75°, field of view (FOV) = 220×220mm, matrix size = 64×64, voxels dimensions = 3.4×3.4×3.4mm, 42 axial oblique slices). T1-weighted structural data were also acquired using a 3D fast spoiled gradient recall brain volume (FSPGR BRAVO) sequence with 186 contiguous sagittal slices (TI = 650ms, bandwidth = 25.0kHz, flip angle = 12°, FOV = 256×256mm, and slice thickness = 1.0mm).

### fMRI Tasks

During the fMRI session, participants performed two different tasks: the Force Concept Inventory (FCI) task and the Physics Knowledge (PK) task.

### FCI Task

Participants completed an in-scanner physics conceptual reasoning task that consisted of questions adapted from the reliable and widely used questionnaire known as the Force Concept Inventory (FCI) (Hestenes et al., 1992; Lasry et al., 2011; Von Korff et al., 2016). The experimental condition presented textual and illustrations of scenarios of objects at rest or in motion and students were asked to choose between a correct Newtonian solution and several reasonable but incorrect non-Newtonian alternatives. Students also completed a sequence of control questions that presented text and figure depictions of everyday physical scenarios that shared similar visual and linguistic characteristics to FCI items (e.g., containing words typically used in introductory Newtonian mechanics, as well as visual presentation and self-paced timing paralleling that of the FCI problems.) Control items, however, tested students on general reading comprehension and/or shape discrimination instead of physics content. Across both FCI and control conditions, questions were presented as blocks composed of three sequential view screens (i.e., “phases”), which consisted of:

- **Phase 1: Scenario:** students viewed text and a figure describing a physical scenario (**Fig. 1A**),
- **Phase II: Question:** students viewed a physics question about the scenario (**Fig. 1B**), and
- **Phase III: Answer:** students responded with their answer out of four possible answer choices (**Fig. 1C**).

**Figure 1.**
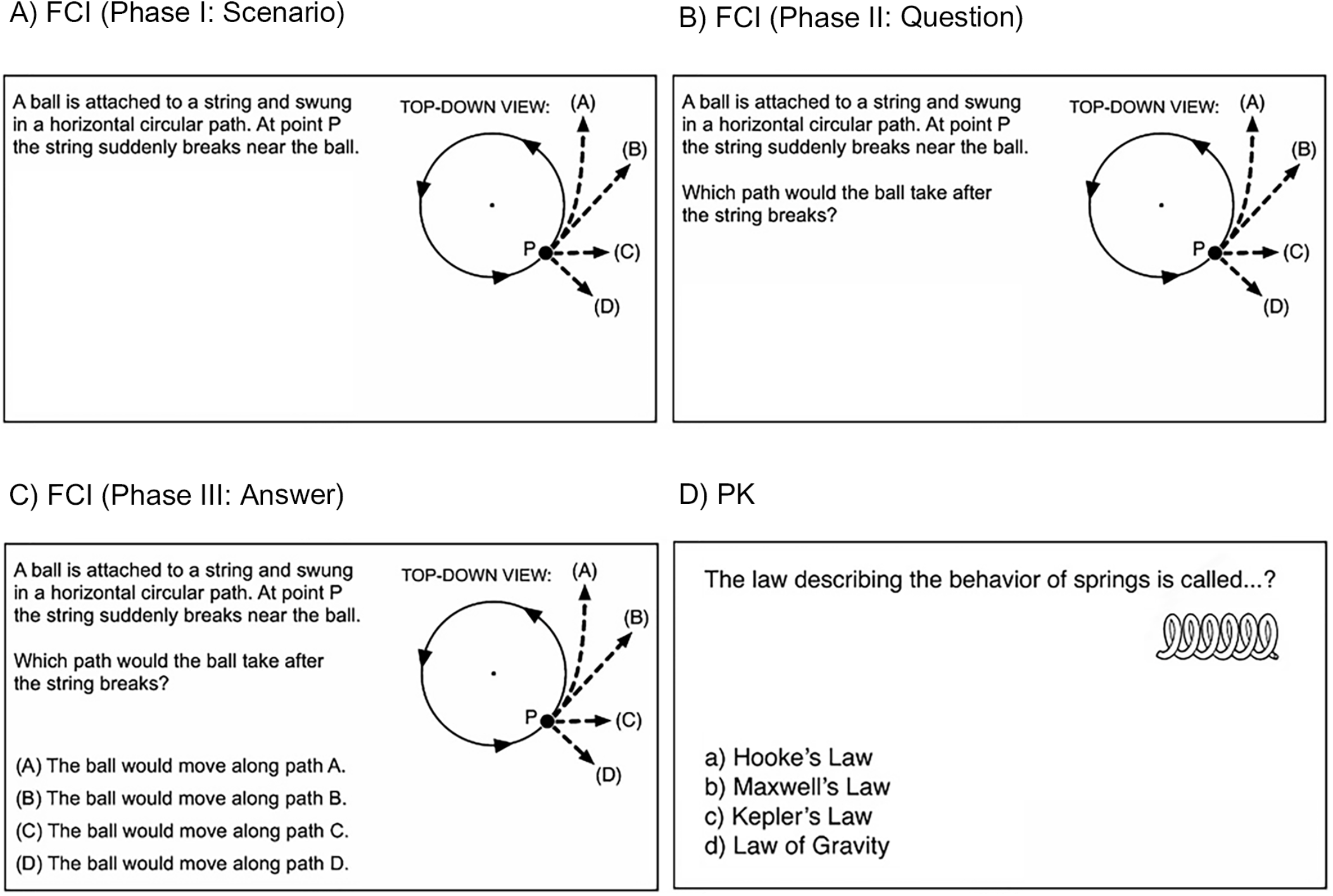
Force Concept Inventory (FCI) and Physics Knowledge (PK) Tasks. Example items of the in-scanner tasks, including the three phases of the FCI task (A) Phase I: Scenario, (B) Phase II: Question, (C) Phase III: Answer and the (B) PK task.

Participants provided a self-paced button press to advance between phases and to provide their final answer. A fixation cross was shown after answer selection and before presentation of the next scenario. FCI and control blocks were of maximum duration 45 sec and were followed by a fixation cross of minimum duration 10 sec. The total duration for each FCI run was 5 min 44 sec; data were collected during three runs for a total duration of ~16 minutes. Control trials were not analyzed in the present study but a description is provided above for completeness. In a previous publication, we demonstrated that our within-scanner version of the FCI task elicited widespread frontoparietal activation of regions of the DAN, VAN, and DMN, and that these activation patterns were elicited differently during each of the three phases (Bartley et al., 2019).

### PK Task

Participants also completed the Physics Knowledge (PK) task. The PK task, adapted from a general knowledge task of semantic retrieval (Elman et al., 2012), was presented in a block-design and probed for brain activation associated with physics-based content knowledge. Students viewed physics questions (e.g., *“What is the value of the acceleration due to gravity on Earth?”*) and corresponding answer choices, such as *“9.81 m/s^2^,15 kg, 10 liters, and 11 ft/s^2^”*) (**Fig. 1D**). A control condition was presented in which students viewed general knowledge questions (e.g., *“What is the tallest mountain in the world?”*) with corresponding answer choices, such as “*Mount Rushmore, Mount Rainier, Mount Everest, or Mount Logan*”). PK and control blocks were 28 seconds long and included four questions per block (6.5 sec per question followed by 0.5 sec of quick fixation). Three blocks of physics or general questions (six question blocks total) were alternated with 10 sec of fixation. The total duration of one run was 4 min 2 sec; data were collected during two runs for a total duration of ~8 minutes. Similar to the FCI task, control trials for the PK task were not analyzed in the present study but are described for completeness.

## Analyses

### Latent Profile Analysis

Latent profile analysis (LPA) is an analytic technique in which participants are assigned (with varying probabilities) into classes (i.e., subpopulations) based on their pattern of responses on a set of indicators. LPA was used in this study to group participants based on STEM-related anxiety profiles (i.e., based on similar self-report of science and math anxiety). LPA was performed in R using the `tidyLPA` package, as well as `mclust`, which uses the Expectation-Maximization algorithm, an approach for maximum likelihood estimation, for model-based clustering and classification (Rosenberg et al., 2018; Scrucca et al., 2016). Default parameters were used in which equal variances across classes and covariances were fixed to 0, which assumes conditional independence of the indicators and that correlation amongst indicators are explained exclusively by the latent classes, and a maximum of four possible classes were specified (Lee et al., 2020). The `compare_solutions` command, available in `tidyLPA`, was used to determine the optimal number of classes by selecting the model with the lowest Bayesian Information Criterion (BIC), entropy, and bootstrapped likelihood ratio test (LRT). BIC is a model selection tool used to select one model from a finite set of possible models and the one with the smallest BIC is considered the “best” candidate (Perrotte et al., 2021). Entropy, which can range from 0 to 1, is an indicator of classification precision and greater entropy suggests that the identified classes are better separated (Lee et al., 2020). Additionally, models with entropy values greater than 0.8 suggest good distinction of the identified classes (Ramaswamy et al., 1993). Bootstrapped LRT produces a p-value for each subsequent model and is an indicator for the degree of fit improvement resulting from adding an additional profile. If the model contains a significant p-value when an additional profile is added, then that suggests the model provides a significant improvement in fit, relative to the previous model with k-1 profiles. Next, `tidyLPA’s plot_profile` command that specifies a 95% confidence interval was used to visualize model classification and assist with interpreting the grouping of the final model. Finally, for the purpose of contextualizing the profiles, we examined the extent to which the profiles differed with respect to age, sex, ethnicity, household income, number of years enrolled at FIU, GPA, generalized anxiety, and accuracy on the FCI and PK tasks (measured as the average number of correct responses).

### fMRI Preprocessing

Each participant’s T1-weighted images were corrected for intensity non-uniformity with `ANT’s N4BiasFieldCorrection` tool (Avants et al., 2008; Tustison et al., 2010). Both anatomical and functional images were preprocessed using `fMRIPrep` (v.1.5.0rc1) (Esteban et al., 2019, 2020). The T1-weighted (T1w) reference, which was used throughout the pipeline, was generated after T1w images were corrected for intensity non-uniformity with ANT’s `N4BiasFieldCorrection. Freesurfer’s mri_robust_template` was used to generate a T1w reference, which was used throughout the entire pipeline (Reuter et al., 2010; Tustison et al., 2010). Nipype’s implementation of `ANT’s antsBrainExtraction` workflow was used to skullstrip the T1w reference using `OASIS30ANTs` as the target template (Gorgolewski et al., 2011). `FSL’s FAST` was used for brain tissue segmentation of the cerebrospinal fluid (CSF), white matter (WM), and gray matter (GM); brain surfaces were reconstructed using `Freesurfer’s recon_all` (Dale et al., 1999; Zhang et al., 2001). Preprocessing of functional images began with selecting a reference volume and generating a skullstripped version using a custom methodology of fMRIPrep. `Freesurfer’s bbregister`, which uses boundary-based registration, was used to coregister the T1w reference to the BOLD reference. The BOLD time series was then resampled onto surfaces of fsaverage5 space and resampled onto their original, native space by applying a single, composite transform to correct for head motion and susceptibility distortions. Additionally, the BOLD time series was high pass filtered, using a discrete cosine filter with a cutoff of 128s (Greve & Fischl, 2009). Several confounding time series were estimated as follows: for each functional run, motion outliers were set at a threshold of 0.5 mm framewise displacement (FD) or 1.5 standardized DVARS. Nuisance signals from the CSF, WM, and whole brain masks were extracted by using a set of physiological regressors, which were extracted to allow for both temporal componentbased noise correction `(tCompCor)` and anatomical component-based noise correction `(aCompCor)` (Behzadi et al., 2007). Additionally, the confound time series derived from head motion estimates were expanded to include its temporal derivatives and quadratic terms, resulting in a total of 24 head motion parameters (i.e., six base motion parameters, six temporal derivatives of six motion parameters, 12 quadratic terms of six motion parameters, and their six temporal derivatives). Estimates for the global, cerebrospinal fluid, and white matter signals were expanded to include their temporal derivatives and quadratic terms, resulting in a total of 12 signal-based parameters (i.e., three base signal parameters, three temporal derivatives of the three base parameters, the three quadratic terms of the base parameters, and the three quadratic terms of the temporal derivatives). Finally, all 24 head motion confound estimates, three high pass filter estimates, and a variable number of `aCompCor` estimates (components that explain the top 50% of the variance) were outputted into a tsv file to be used for later denoising steps (Satterthwaite et al., 2013).

### Parcellation and Task-Based Connectivity Analyses

Additional data analysis was conducted in `IDConn`, a pipeline that bundles several commonly used neuroimaging software packages to create workflows examining functional brain connectivity (Bottenhorn & Salo, 2022). Each participant’s preprocessed FCI and PK task-based fMRI data were parcellated according to a functionally derived, whole-brain parcellation. Network-level identification of the DAN, VAN, and DMN was carried out using the 17-network parcellation developed by Yeo et al. (2011), using individual nodes within the networks as identified by Kong et al. (2021) (**Fig. 2**). Confounding time series identified by `fMRIPrep`, along with the six head motion estimates from `FSL’s MCFLIRT` and outlier volumes (FD > 0.5 mm or 1.5 standardized DVARS) identified during preprocessing, were regressed out during analysis in `IDConn`. Each functional task time series was standardized and the average per-network time series were extracted for each participant and task condition (averaged across runs), allowing assessment of between-network connectivity. Similarly, the average per-node time series was extracted for each participant and task condition (averaged across runs), allowing assessment of within-network connectivity. Importantly, given the relatively long duration of FCI trials (i.e., 45 sec), we separately extracted average network- and node-level time series for FCI Phase I (Scenario), Phase II (Question), and Phase III (Answer). Adjacency matrices were constructed per participant, per functional task using `Nilearn` (v. 0.3.1, http://nilearn.github.io/index.html), a Python (v. 2.7.13) module, built on `scikit-learn`, for the statistical analysis of neuroimaging data, by computing the pairwise Pearson’s correlations between each pair of regions, resulting in a 400×400 region-wise correlation matrix for each participant per condition per task (Bottenhorn et al., 2021; Medaglia, 2017). From these matrices, we assessed the between-network connectivity for DAN-VAN, DAN-DMN, and VAN-DMN, as well as the within-network connectivity for the DAN, VAN, and DMN. Between- and within-network connectivity were assessed for each participant and for both the FCI and PK tasks.

**Figure 2.**
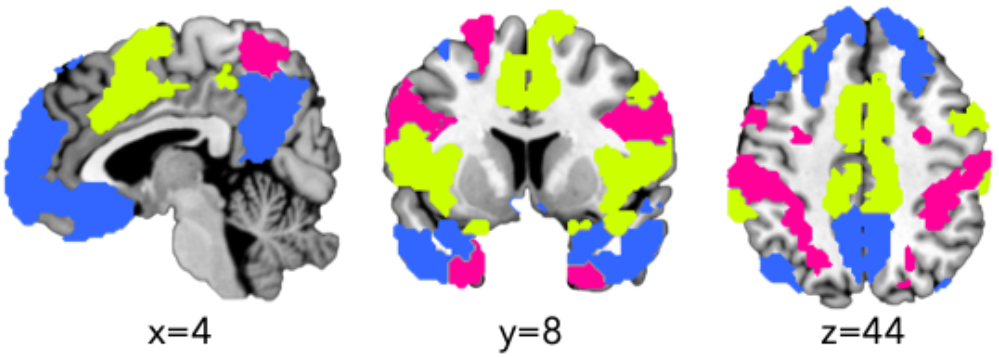
DAN, VAN, and DMN Network Parcellation. Each participant’s preprocessed fMRI data were parcellated using the 17-network Yeo et al. (2011) parcellation to identify the dorsal attention network (DAN; pink); ventral attention (VAN; yellow), and default mode network (DMN; blue).

### Statistical Analyses

Statistical modeling was conducted with the Lavaan package (Rosseel, 2012), which is available in R. Regression models were generated to evaluate between-network connectivity between DAN-VAN, DAN-DMN, and VAN-DMN during both the FCI and PK tasks. In addition, regression models were also generated to evaluate within-network connectivity for the DAN, VAN, and DMN during the FCI and PK tasks. The FCI models included three observed variables for each phase, including the average between- or within-network connectivity values during Phase I (Scenario), Phase II (Question), and Phase III (Answer). The PK models included a single observed variable, which was the average between- or within-network connectivity values for each of the PK or control conditions. For each model, the residual variance and intercept for the observed variable were specified. The main explanatory variable of interest was LPA-based class assignment. Age, sex, ethnicity, household income, number of years enrolled at FIU, GPA, and generalized anxiety were included as covariates.

## Results

### Latent Profile Analysis

**Table 2** presents the model fit indices for the four LPA models of science and math anxiety. The 4-profile model had the lowest Bayesian Information Criteria (BIC = 615.33), which suggested that this model, relative to the other three, demonstrated the greatest improvement in fit. This interpretation was supported by the results of the bootstrapped LRT p-values, which showed that the differences in improvement of fit between the 1- and 2-profile models (p = 0.01), the 2- and 3-profile models (p = 0.01), and the 3- and 4-profile models (p = 0.01) were all significant. Furthermore, the 4-profile model had an entropy value greater than 0.8, which suggested a good separation of the identified classes (Ramaswamy et al., 1993). Thus, the 4-profile model was selected based on BIC, entropy, bootstrapped LRT, and interpretability of classes.

**Table 2.** Latent Profile Analysis Model Comparisons.

| # of Profiles | BIC | Entropy | Bootstrapped LRT *p*-values |
| --- | --- | --- | --- |
| 1 | 715.36 | 1.00 | - |
| 2 | 663.07 | 0.91 | 0.01 |
| 3 | 629.27 | 0.89 | 0.01 |
| 4 | 615.33 | 0.88 | 0.01 |

*Note.* BIC = Bayesian Information Criteria, LRT = Lo-Mendell-Rubin Adjusted Likelihood Ratio Test.

**Table 3** presents the mean z-scores for science and math anxiety across the four profiles. The `tidyLPA get_estimates` command was used to determine if the mean science and anxiety for each profile was significant based on a p-value < 0.05. The first profile represented 6.5% of the sample (n = 8) and was labeled as High STEM Anxiety as both science and math anxiety were significantly above zero. The second profile represented 59.3% of the sample (n = 73) and was labeled as Low STEM Anxiety as both science and math anxiety means were significantly below zero. The third profile represented 21.9% of the sample (n = 27) and was labeled as High Math Anxiety as only math anxiety was significantly above zero. The fourth profile represented 4.1% of the sample (n = 5) and was labeled as High Science Anxiety as only science anxiety was significantly above zero.

**Table 3.** Parameter Estimates for Science and Math Anxiety.

| Profile | Science Anxiety |  |  | Math Anxiety |  |  |
| --- | --- | --- | --- | --- | --- | --- |
| | $\mu$ | $\sigma^2$ | $p$-value | $\mu$ | $\sigma^2$ | $p$-value |
| High STEM Anxiety (HSA) | 1.490 | 0.212 | <0.001 | 2.220 | 0.300 | <0.001 |
| Low STEM Anxiety (LSA) | -0.499 | 0.212 | <0.001 | -0.572 | 0.300 | <0.001 |
| High Math Anxiety (HMA) | 0.280 | 0.212 | 0.088 | 0.776 | 0.300 | <0.001 |
| High Science Anxiety (HSA) | 3.330 | 0.212 | <0.001 | 0.016 | 0.300 | 0.970 |

Note. Profile means and variances are represented by $\mu$ and $\sigma^2$ , respectively.

To ensure clearly defined class membership, we restricted assignment to profiles to those whose posterior probabilities were 0.70 or higher. Of the 123 total participants, 100 of participants (81.3%) had posterior probabilities greater than 0.70. This included 73 Low STEM Anxiety and 27 High Math Anxiety participants. Given low sample sizes for the High STEM Anxiety and High Science Anxiety groups, these profiles were excluded from subsequent analysis. Thus, further analysis only focused on examining differences between the High Math Anxiety and Low STEM Anxiety groups. **Fig. 3** illustrates the z-scores for science and math anxiety across these two profiles.

**Figure 3.**
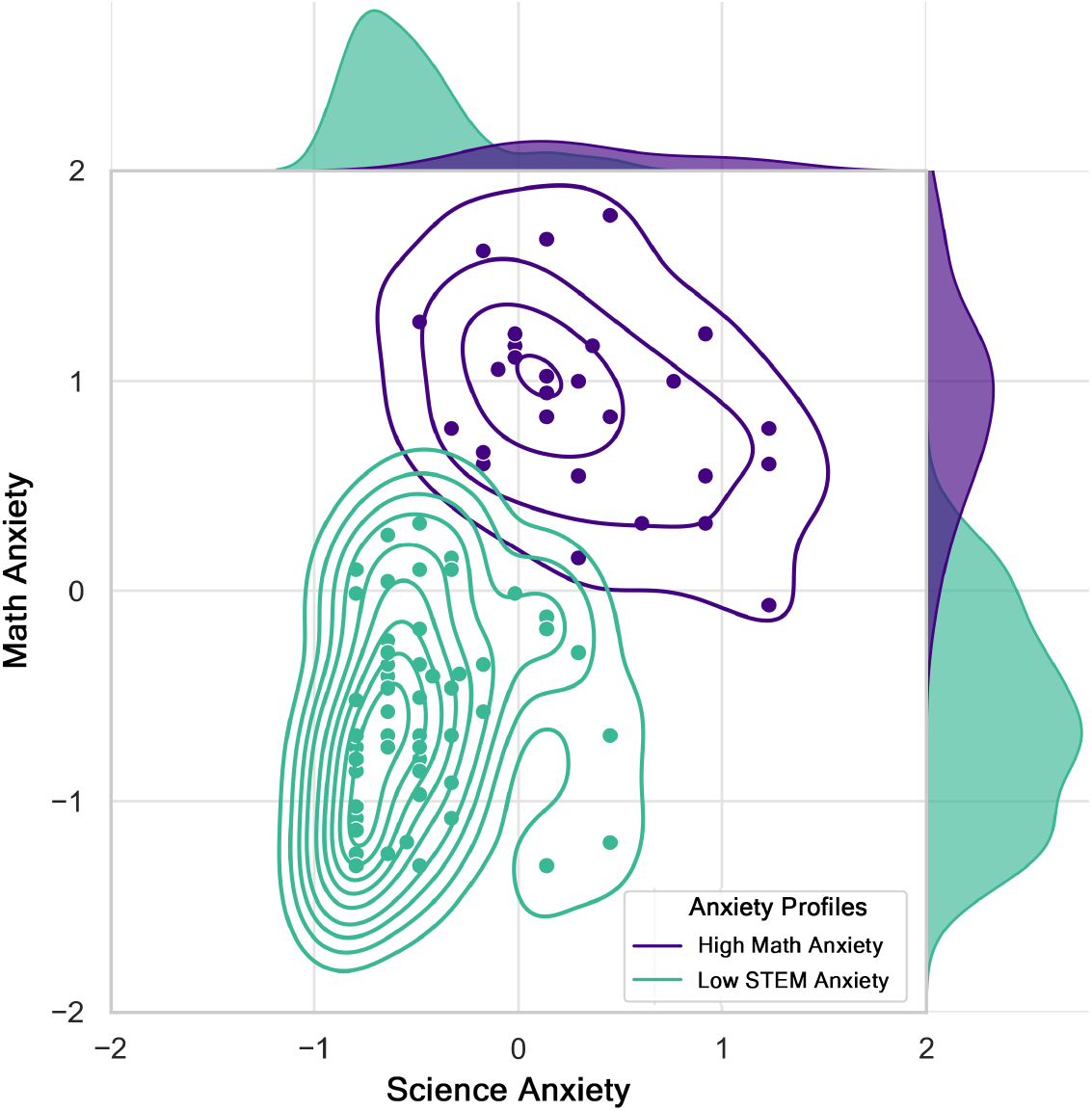
Science and Math Anxiety Scores for High Math Anxiety and Low STEM Anxiety Groups. A joint kernel density estimate plot showing the distributions of standardized math and science anxiety scores for High Math Anxiety (purple) and Low STEM Anxiety (green) students.

### Demographic Differences Across Profiles

Next, we explored demographic differences across the High Math Anxiety and Low STEM Anxiety profiles. Results from chi-square tests of association indicated that we could not reject the null hypothesis of no difference on the basis of sex (χ^2^ = 3.361, df = 1, p = 0.067), ethnicity (0.0870, df = 1, p = 0.768), household income (χ^2^= 2.368, df = NA, p = 0.809), or number of years enrolled at FIU (χ^2^= 4.501, df = NA, p = 0.219). Furthermore, results from t-tests indicated that we could not reject the null hypothesis of no difference in terms of average age (t = 1.511, df = 48.927, p = 0.137) or GPA (t = −1.132, df = 67.728, p = 0.262). Importantly, and contrary to our hypotheses, we could not reject the null hypothesis on fMRI task performance between groups, with no significant differences in terms of FCI accuracy (t = 1.221, df = 60.613, p = 0.227), or PK accuracy (t = −1.128, df = 49.695, p = 0.265). Lastly, High Math Anxiety students exhibited significantly increased generalized anxiety compared to Low STEM Anxiety students (t = 2.481, df = 31.732, p = 0.0186).

### Profile Membership Effects: Between-Network Connectivity

We examined whether there were significant differences between High Math Anxiety and Low STEM Anxiety students in terms of between-network connectivity for the DAN, VAN, and DMN. **Table 4** presents the between-network connectivity differences during the FCI task. Results indicated no significant differences in connectivity during FCI Phase I (Scenario) or Phase II (Question). However, during Phase III (Answer), High Math Anxiety students exhibited significantly reduced between-network connectivity (i.e., DAN-VAN, VAN-DMN, and DAN-DMN) relative to Low STEM Anxiety students. Distributions of between-network connectivity values for Phase III of the FCI task are displayed in **Fig. 4A**. Student age, sex, ethnicity, household income, number of years enrolled at FIU, GPA, and generalized anxiety did not significantly explain variation in between-network connectivity across all FCI phases.

**Table 4.**
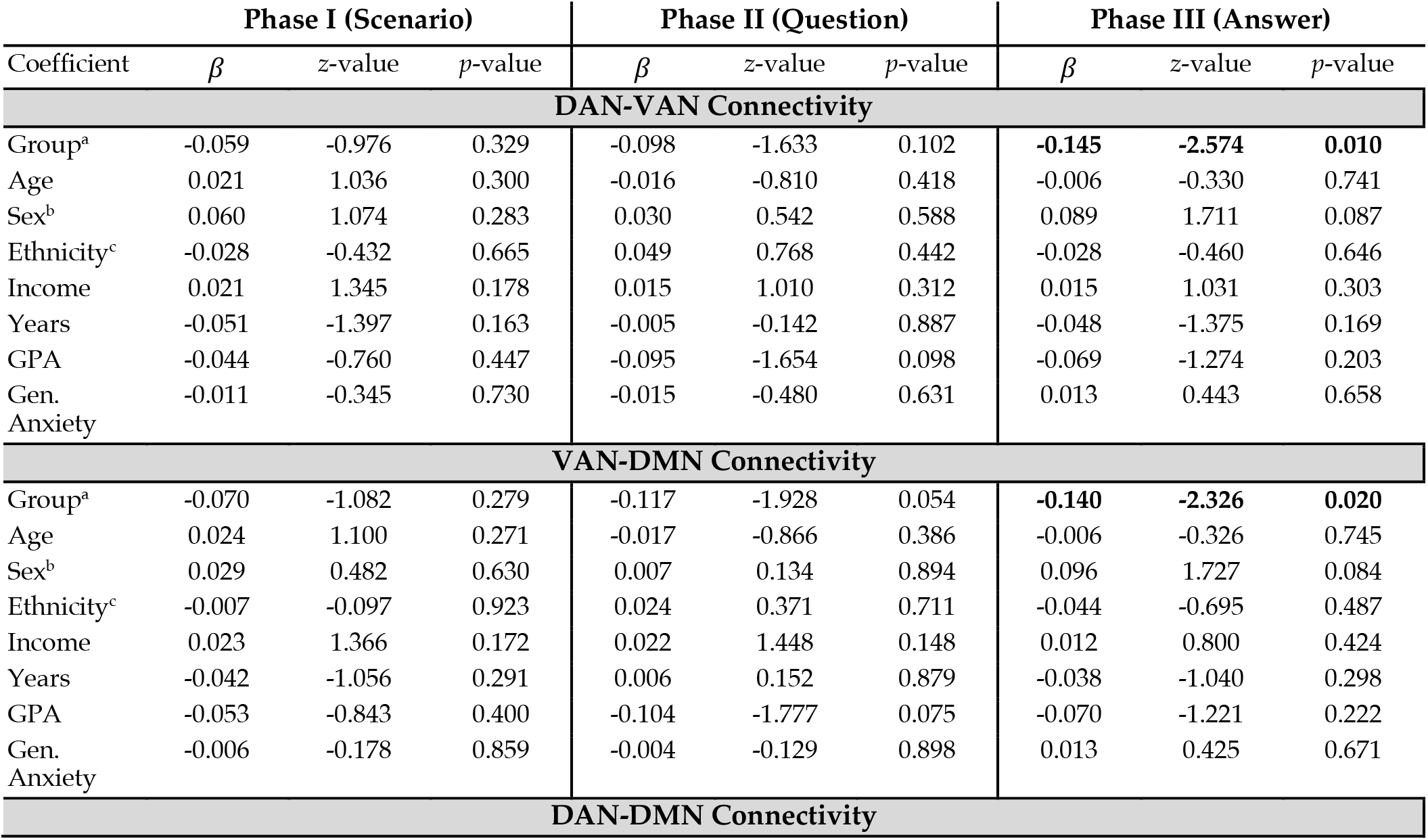
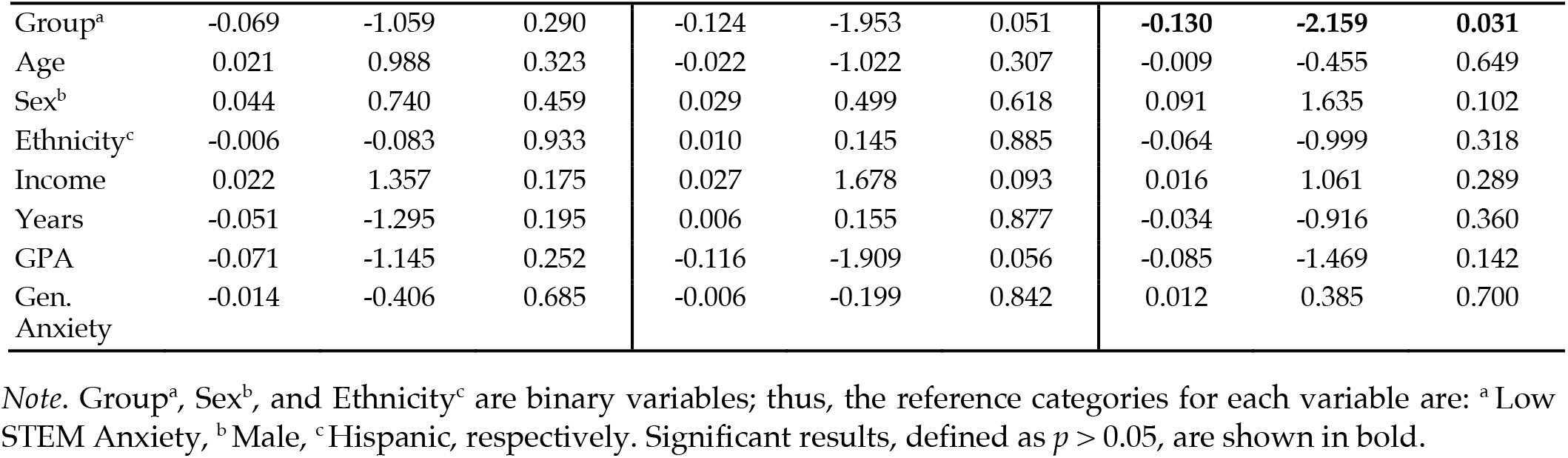
Between-Network Connectivity during the Force Concept Inventory (FCI) Task.

**Figure 4.**
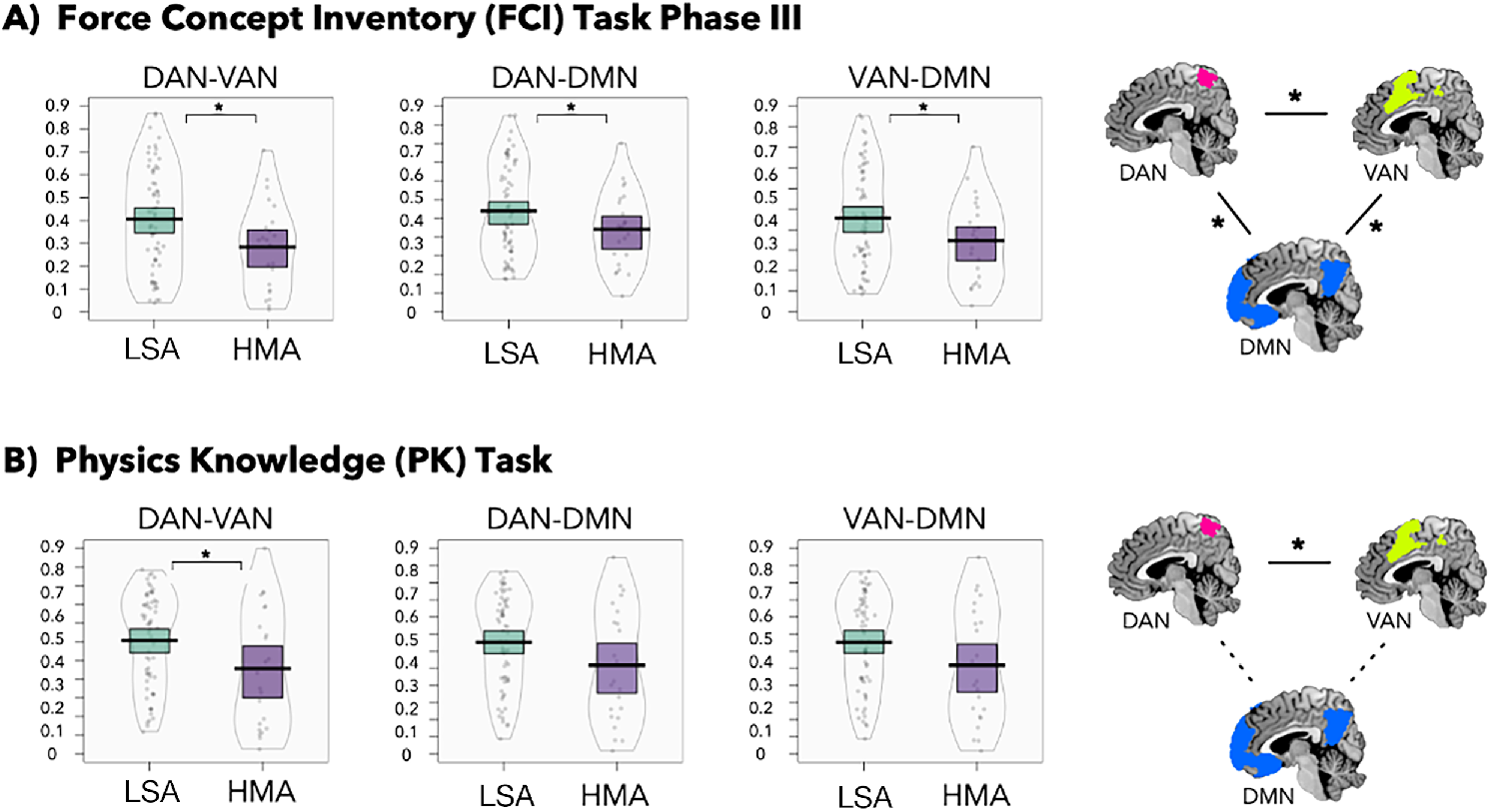
Between-Network Connectivity Results. Distributions of between-network connectivity values during the A) Force Concept Inventory (FCI) Task Phase III and B) physics knowledge (PK) task among Low STEM Anxiety (LSA; green) and High Math Anxiety (HMA; purple) students. Pirate plots with asterisks denote significant differences between groups. For each task, the observed between-network differences are illustrated with DAN, VAN, and DMN topographical visualization. Asterisks accompanied by a solid line denote significant differences between groups; dotted lines represent no significant group differences.

**Table 5** presents the between-network connectivity differences during the PK task. Results indicated no significant differences in DMN-related between-network connectivity (i.e., VAN-DMN and DAN-DMN). However, High Math Anxiety students exhibited significantly reduced DAN-VAN connectivity during the PK task relative to Low STEM Anxiety students. Distributions of between-network connectivity values for the PK task are displayed in **Fig. 4B**. As with the FCI task, age, sex, ethnicity, household income, number of years enrolled at FIU, GPA, and generalized anxiety were not significant predictors for between-network connectivity for the PK task.

**Table 5.** Between-Network Connectivity during Physics Knowledge (PK) Task.

| Coefficient | $\beta$ | z-value | p-value | $\beta$ | z-value | p-value | $\beta$ | z-value | p-value |
| --- | --- | --- | --- | --- | --- | --- | --- | --- | --- |
|  | DAN-VAN Connectivity |  |  | VAN-DMN Connectivity |  |  | DAN-DMN Connectivity |  |  |
| Group<sup>a</sup> | **-0.150** | **-2.005** | **0.045** | -0.104 | -1.347 | 0.178 | -0.155 | -1.923 | 0.054 |
| Age | -0.007 | -0.300 | 0.764 | -0.007 | -0.282 | 0.778 | -0.001 | -0.047 | 0.962 |
| Sex<sup>b</sup> | -0.016 | -0.235 | 0.814 | 0.002 | 0.032 | 0.975 | 0.019 | 0.252 | 0.801 |
| Ethnicity<sup>c</sup> | 0.014 | 0.177 | 0.859 | -0.002 | -0.027 | 0.978 | -0.002 | -0.028 | 0.978 |
| Income | -0.019 | -0.993 | 0.321 | -0.026 | -1.324 | 0.185 | -0.021 | -1.007 | 0.314 |
| Years | 0.023 | 0.505 | 0.613 | 0.023 | 0.483 | 0.629 | 0.014 | 0.276 | 0.783 |
| GPA | -0.063 | -0.883 | 0.377 | -0.094 | -1.256 | 0.209 | -0.077 | -0.999 | 0.318 |
| Gen. Anxiety | -0.015 | -0.393 | 0.694 | -0.014 | -0.362 | 0.718 | -0.001 | -0.013 | 0.989 |

Note. Group<sup>a</sup>, Sex<sup>b</sup>, and Ethnicity<sup>c</sup> are binary variables; thus, the reference categories for each variable are: <sup>a</sup> Low STEM Anxiety, <sup>b</sup> Male, <sup>c</sup> Hispanic, respectively. Significant results, defined as $p > 0.05$ , are shown in bold.

### Profile Membership Effects: Within-Network Connectivity

Lastly, we examined whether there were significant differences between High Math Anxiety and Low STEM Anxiety students in terms of within-network connectivity for the DAN, VAN, and DMN. **Table 6** presents the with-network connectivity differences during the FCI task. Results indicated no differences in connectivity between groups during FCI Phase I (Scenario) or Phase II (Question). However, during Phase III (Answer), High Math Anxiety students exhibited significantly reduced within-network connectivity (i.e., DAN, VAN, and DMN) relative to Low STEM Anxiety students. Distributions of within-network connectivity values for Phase III of the FCI task are displayed in **Fig. 5A**. Student age, sex, ethnicity, household income, number of years enrolled at FIU, GPA, and generalized anxiety did not significantly explain variation in within-network connectivity across all FCI phases.

**Table 6.** Within-Network Connectivity during Force Concept Inventory (FCI) Task.

| Coefficient | Phase I (Scenario) |  |  | Phase II (Question) |  |  | Phase III (Answer) |  |  |
| --- | --- | --- | --- | --- | --- | --- | --- | --- | --- |
| | $\beta$ | z-value | p-value | $\beta$ | z-value | p-value | $\beta$ | z-value | p-value |
| **Within-DAN Connectivity** |  |  |  |  |  |  |  |  |  |
| Group<sup>a</sup> | -0.052 | -0.893 | 0.372 | -0.105 | -1.818 | 0.069 | **-0.141** | **-2.525** | **0.012** |
| Age | 0.018 | 0.947 | 0.344 | -0.020 | -1.060 | 0.289 | -0.010 | -0.536 | 0.592 |
| Sex<sup>b</sup> | 0.046 | 0.846 | 0.398 | 0.008 | 0.156 | 0.876 | 0.089 | 1.719 | 0.086 |
| Ethnicity<sup>c</sup> | -0.027 | -0.426 | 0.670 | 0.059 | 0.958 | 0.338 | -0.025 | -0.417 | 0.676 |
| Income | 0.017 | 1.137 | 0.256 | 0.020 | 1.333 | 0.182 | 0.014 | 0.988 | 0.323 |
| Years | -0.054 | -1.510 | 0.131 | 0.004 | 0.112 | 0.911 | -0.033 | -0.973 | 0.330 |
| GPA | -0.040 | -0.704 | 0.482 | -0.076 | -1.370 | 0.171 | -0.065 | -1.210 | 0.226 |
| Gen. Anxiety | -0.004 | -0.138 | 0.891 | 0.001 | 0.029 | 0.977 | 0.020 | 0.698 | 0.485 |
| **Within-VAN Connectivity** |  |  |  |  |  |  |  |  |  |
| Group<sup>a</sup> | -0.049 | -0.843 | 0.399 | -0.055 | -0.957 | 0.339 | **-0.142** | **-2.540** | **0.011** |
| Age | 0.019 | 0.974 | 0.330 | -0.019 | -1.022 | 0.307 | -0.005 | -0.263 | 0.792 |
| Sex<sup>b</sup> | 0.046 | 0.849 | 0.396 | 0.019 | 0.356 | 0.722 | 0.090 | 1.742 | 0.082 |
| Ethnicity<sup>c</sup> | -0.024 | -0.385 | 0.700 | 0.058 | 0.950 | 0.342 | -0.026 | -0.431 | 0.666 |
| Income | 0.015 | 1.001 | 0.317 | 0.012 | 0.819 | 0.413 | 0.012 | 0.862 | 0.389 |
| Years | -0.040 | -1.105 | 0.269 | 0.011 | 0.309 | 0.758 | -0.040 | -1.168 | 0.243 |
| GPA | -0.036 | -0.646 | 0.518 | -0.076 | -1.390 | 0.164 | -0.049 | -0.920 | 0.357 |
| Gen. Anxiety | -0.006 | -0.214 | 0.830 | -0.017 | -0.593 | 0.553 | 0.014 | 0.476 | 0.634 |
| **Within-DMN Connectivity** |  |  |  |  |  |  |  |  |  |
| Group<sup>a</sup> | -0.058 | -1.009 | 0.313 | -0.084 | -1.527 | 0.127 | **-0.129** | **-2.398** | **0.016** |
| Age | 0.022 | 1.122 | 0.262 | -0.020 | -1.091 | 0.275 | 0.000 | 0.008 | 0.994 |
| Sex<sup>b</sup> | 0.005 | 0.095 | 0.924 | -0.015 | -0.294 | 0.769 | 0.086 | 1.735 | 0.083 |
| Ethnicity<sup>c</sup> | -0.006 | -0.099 | 0.921 | 0.069 | 1.174 | 0.240 | -0.049 | -0.851 | 0.395 |
| Income | 0.027 | 1.873 | 0.061 | 0.019 | 1.353 | 0.176 | 0.015 | 1.126 | 0.260 |
| Years | -0.043 | -1.207 | 0.227 | -0.010 | -0.304 | 0.761 | -0.038 | -1.151 | 0.250 |
| GPA | -0.048 | -0.868 | 0.385 | -0.080 | -1.515 | 0.130 | -0.073 | -1.422 | 0.155 |
| Gen. Anxiety | -0.001 | -0.048 | 0.962 | -0.009 | -0.329 | 0.742 | 0.004 | 0.157 | 0.875 |

Note. Group<sup>a</sup>, Sex<sup>b</sup>, and Ethnicity<sup>c</sup> are binary variables; thus, the reference categories for each variable are: <sup>a</sup> Low STEM Anxiety, <sup>b</sup> Male, <sup>c</sup> Hispanic, respectively. Significant results, defined as $p > 0.05$ , are shown in bold.

**Figure 5.**
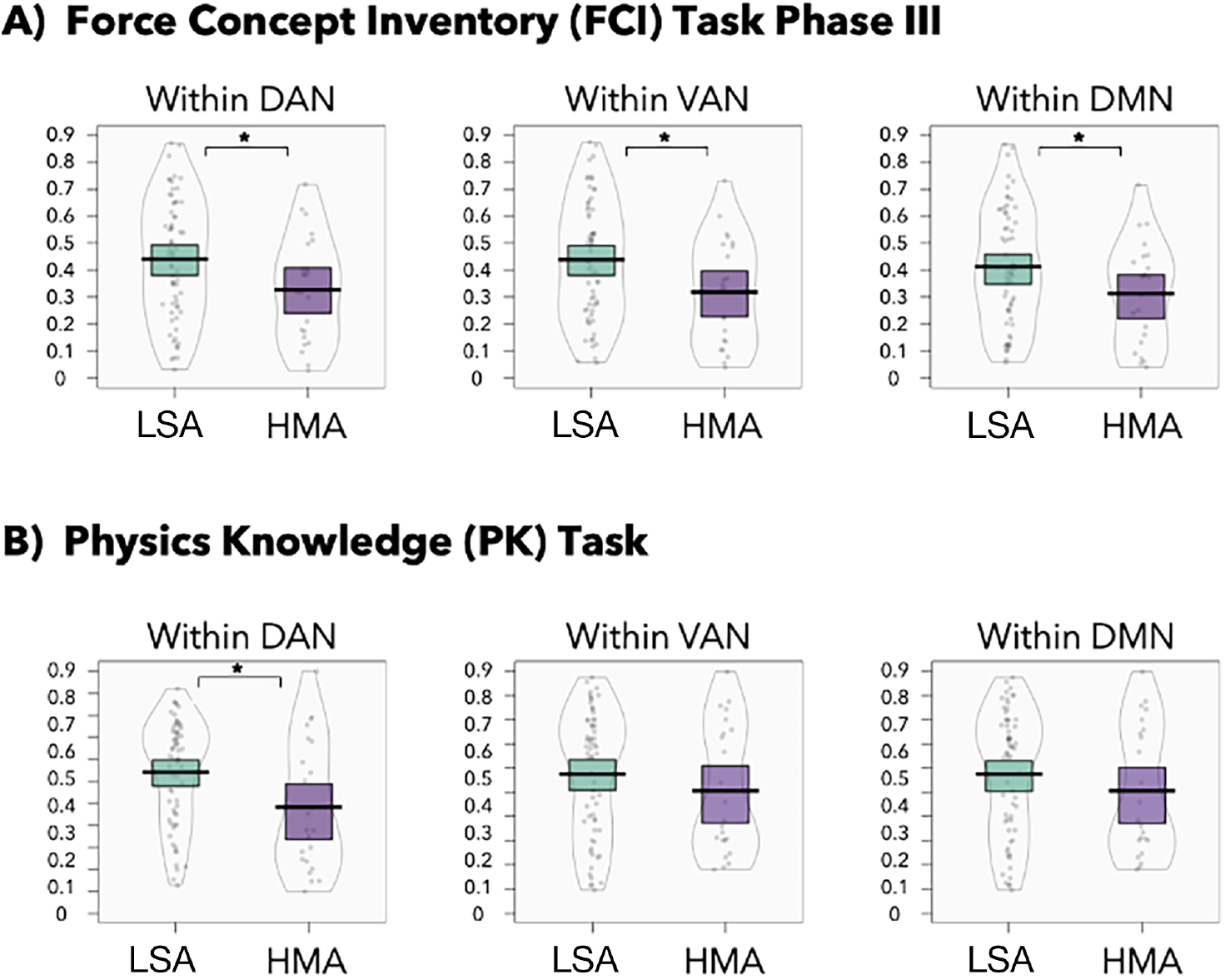
Within-Network Connectivity Results. Distributions of within-network connectivity values during the A) Force Concept Inventory (FCI) Task Phase III and B) physics knowledge (PK) task among Low STEM Anxiety (LSA; green) and High Math Anxiety (HMA; purple) students. Pirate plots with asterisks denote significant differences between groups.

**Table 7** presents the within-network connectivity differences during the PK task. Results indicated no significant differences in within-network connectivity between groups for the VAN or DMN. However, High Math Anxiety students exhibited significantly reduced within-DAN connectivity during the PK task relative to Low STEM Anxiety students. Distributions of within-network connectivity values for the PK task are displayed in **Fig. 5B**. As with the FCI task, age, sex, ethnicity, household income, number of years enrolled at FIU, GPA, and generalized anxiety were not significant predictors for within-network connectivity for the PK task.

**Table 7.** Within-Network Connectivity during Physics Knowledge (PK) Task.

| Coefficient | $\beta$ | z-value | p-value | $\beta$ | z-value | p-value | $\beta$ | z-value | p-value |
| --- | --- | --- | --- | --- | --- | --- | --- | --- | --- |
|  | **DAN Within-Network Connectivity** |  |  | **VAN Within-Network Connectivity** |  |  | **DMN Within-Network Connectivity** |  |  |
| Group<sup>a</sup> | **-0.165** | **-2.386** | **0.017** | -0.122 | -1.645 | 0.100 | -0.086 | -1.213 | 0.225 |
| Age | -0.005 | -0.200 | 0.841 | -0.010 | -0.386 | 0.699 | -0.000 | -0.006 | 0.995 |
| Sex<sup>b</sup> | -0.030 | -0.467 | 0.641 | 0.009 | 0.129 | 0.897 | -0.005 | -0.072 | 0.943 |
| Ethnicity<sup>c</sup> | 0.046 | 0.630 | 0.529 | 0.001 | 0.017 | 0.986 | 0.004 | 0.060 | 0.952 |
| Income | -0.019 | -1.102 | 0.271 | -0.021 | -1.121 | 0.262 | -0.029 | -1.630 | 0.103 |
| Years | 0.017 | 0.399 | 0.690 | 0.018 | 0.395 | 0.693 | 0.015 | 0.344 | 0.731 |
| GPA | -0.062 | -0.934 | 0.350 | -0.067 | -0.945 | 0.345 | -0.086 | -1.267 | 0.205 |
| Gen. Anxiety | 0.004 | -0.111 | 0.912 | -0.029 | -0.763 | 0.445 | -0.002 | -0.051 | 0.960 |

Note. Group<sup>a</sup>, Sex<sup>b</sup>, and Ethnicity<sup>c</sup> are binary variables; thus, the reference categories for each variable are: <sup>a</sup> Low STEM Anxiety, <sup>b</sup> Male, <sup>c</sup> Hispanic, respectively. Significant results, defined as $p > 0.05$ , are shown in bold.

## Discussion

The current study examined associations between STEM-related anxiety and between- and within-network connectivity for the DAN, VAN, and DMN among university physics students. LPA conducted on science and math anxiety scores identified four student profiles; however only two profiles, which included High Math Anxiety and Low STEM Anxiety students, were examined due to low sample size. No significant differences in within- and between-network connectivity were observed between these two profiles in terms of student age, sex, household income, ethnicity, years of education, GPA, or physics task performance. Significant differences in within- and between-network connectivity were observed between profiles in terms of their generalized anxiety, with High Math Anxiety students exhibiting increased generalized anxiety compared to Low STEM Anxiety students. Group differences in between- and within-network connectivity for the DAN, VAN, and DMN were examined during the FCI and PK tasks, which measure physics-based conceptual reasoning and content knowledge, respectively. Results demonstrated no significant group differences in task-based connectivity during FCI Phase I (Scenario) and Phase II (Question). However, during FCI Phase III (Answer), High Math Anxiety students exhibited significantly reduced between-network (i.e., DAN-VAN, VAN-DMN, and DAN-DMN) connectivity and within-network (i.e., DAN, VAN, and DMN) connectivity compared to Low STEM Anxiety students. Regarding the PK task, no significant group differences were observed in DMN-related between-network connectivity (i.e., VAN-DMN and DAN-DMN) or within-network connectivity for the VAN and DMN. However, during the PK task, High Math Anxiety students exhibited significantly reduced DAN-VAN and within-DAN connectivity compared to Low STEM Anxiety students. These outcomes provide insight into how between- and within-network connections (i.e., DAN, VAN, and DMN) are altered among those elevated and reduced STEM-related anxiety.

### Physics Cognition During the FCI and PK Tasks

The current study used two different fMRI tasks to disentangle attentional control processes to further understanding of the role of STEM-related anxiety on physics-based cognition. During both the FCI and the PK tasks, students responded to physics-based questions. A key difference between tasks is their relative duration; the PK task has a relatively short mean response time of (4.3 sec), while Phase III of the FCI has a much longer mean response time of 20.2 sec. Thus, while both tasks rely on engagement of the DAN for attentional maintenance and control, the FCI also engages the DMN, particularly during Phase III (Answer), likely due to mental exploration and sustained cognition needed to generate their answers (Bartley et al., 2019). We found significant and widespread group differences in task-based connectivity within and between the DAN, VAN, and DMN during Phase III of the FCI, but no differences during Phases I or II, suggesting that attentional control processes are highly relevant and confirm the ACT framework during physics conceptual reasoning and answer selection, but not problem initiation or question presentation. These results are broadly consistent with findings from prior studies investigating the role of DMN-related processing associated with STEM-related anxiety. It has previously been shown that individuals with elevated STEM-related anxiety exhibit disrupted DMN-related functioning (Qiao et al., 2020), have diffuse and unstructured network connectivity (Klados et al., 2019), and experience increased levels of threat avoidance and rumination that may result in deficits in attentional control (Pizzie & Kraemer, 2017), compared to their less-anxious peers. Such results are consistent with ACT since they suggest that highly anxious individuals may utilize more cognitive resources to sustain DMN activity. Consequently, highly anxious individuals may have difficulty in suppressing interference caused by negative emotional information and may experience less mental flexibility in shifting their attention from an internal introspective state to external environmental stimuli, which may lead to performance deficits in tasks requiring high cognitive demands (Qiao et al., 2020). Furthermore, increased ruminators exhibit decreased DMN-related connectivity (i.e., both within-DMN and DAN-DMN) associated with distractor inhibition, emotional regulation, and attentional control (Liu et al., 2020; Roberts et al., 2021; Rosenbaum et al., 2018). In addition, DMN-related connectivity has been linked to distractor suppression (Poole et al., 2016). Together, these findings may offer insight into our significant findings during FCI Phase III and DMN engagement during this task stage may potentially reflect increased rumination among students with elevated STEM anxiety.

In contrast to the FCI results, we found no DMN-related connectivity group differences during the PK task. Significant group differences during the PK task were limited to the DAN, including DAN-VAN and within-DAN connectivity. These results highlight the different cognitive processes and systems at play during the PK task compared to the FCI task. While both the FCI and PK tasks require engagement of DMN regions for successful task execution and retrieval of physics-based content knowledge, our results indicate that it is the dynamics of the DAN that are especially salient in the context of performing the PK task among students with elevated STEM anxiety. The DAN is known to be integral for attention and is not considered a memory system per se (Lückmann et al., 2014), but prior work has demonstrated that top-down attentional control in the DAN plays an important role for successful episodic retrieval when retrieval of specific perceptual information or details is required (Guerin et al., 2012) or during remembrance of external stimuli (Stawarczyk et al., 2018). During the PK task, attentional control is critically important as participants attend to visual cues intended to reactivate systems that encode the material (Lückmann et al., 2014) and trigger recollection of physics-based knowledge, drawing on working memory processes to retrieve information from long-term memory (Fukuda & Woodman, 2017). Such attentional control requires appropriate coordination between the DAN and VAN for attention maintenance and reorienting attention to salient stimuli, respectively (Boon et al., 2020; Corbetta & Shulman, 2002; M. Eysenck et al., 2005).

### Attentional Control Theory and Task Performance

A substantial body of work has previously demonstrated that DMN- and DAN-related connectivity is associated with reduced task performance. During cognitively demanding tasks, greater positive correlation between DAN and DMN activity typically results in greater variability in task performance (Anticevic et al., 2012; Kelly et al., 2008). Konishi et al. (2015) observed that activation in regions associated with working memory was accompanied by significant transient activation in the medial prefrontal cortex (mPFC) and posterior cingulate cortex (PCC), both core hubs of the DMN, for correct responses. Esterman et al. (2013) found that fluctuations in DMN activity had both beneficial and detrimental effects on performance during a sustained attention task. Specifically, while moderate DMN activity corresponded to periods of reduced variability in response time and less error-prone performance, extreme peaks in DMN engagement during these periods predicted lapses in sustained attention and preceded errors. Additionally, positive coupling between the DMN and right inferior frontal gyrus increases speed and accuracy in detecting task-relevant features, which is an important functional component of goal-directed tasks (Elton & Gao, 2015). Furthermore, positive correlations between the mPFC and PCC and the right anterior insula has been associated with performance deficits during sustained attention tasks (Fortenbaugh et al., 2017; Kucyi et al., 2016). Within- and between-network connectivity have also played a role in distractor suppression. Elevated within-network DMN connectivity is predictive of increased distractor suppression, which corresponds to better task performance; however, DMN hyperconnectivity to both the DAN and VAN is predictive of poorer distraction suppression, leading to worse task performance (Poole et al., 2016).

Given these prior results, it was surprising that the observed differences in DMN- and DAN-related connectivity were not linked to FCI and PK differences in task performance between High Math Anxiety and Low STEM Anxiety students. While some previous studies observed reduced task performance among participants with elevated anxiety (Basanovic et al., 2022; Wong et al., 2013; Vytal et al., 2012), other studies noted no associations between anxiety and task performance (Aylward et al., 2017; Derakshan et al., 2009). Importantly, the current lack of group differences in task performance remains consistent within the ACT framework (Eysenck et al., 2007). The emphasis of ACT is on the potential impairment of efficient goal-directed attentional control processing due to the increased probability of cognitive resources being diverted from the task to process anxiety-related stimuli. ACT thus predicts that impaired processing efficiency may manifest as reduced task performance if sufficient auxiliary cognitive resources are not available to maintain performance effectiveness at the cost of impaired efficiency (Eysenck et al., 2007). Consequently, anxious individuals may avoid decrements in task performance through compensatory mechanisms, such as increased cognitive effort and use of cognitive resources (Barker et al., 2018; Eysenck et al., 2007; Minnick et al., 2020). It is possible that anxious participants in this study did not experience elevated anxiety to the degree in which it had a noticeable effect on task performance (Kim et al., 2021); it may be useful to consider adding cognitive load manipulations to future physics-related fMRI tasks.

### Limitations

This study is characterized by several limitations. First, this study is limited by its sample size, which may not have been large enough to i) assign sufficient participants to the two STEM-anxiety profiles that weren’t examined, which included the High STEM anxiety and High Science Anxiety profiles, and ii) detect additional STEM-anxiety related student profiles (Spurk et al., 2020). Our study included 123 student participants and LPA yielded four profiles of students of imbalanced group sizes. As a result, two of those groups included less than 10.6% of the total sample and were excluded from the subsequent connectivity analyses due to power concerns. The present study thus focused only on group differences in task-based connectivity among High Math Anxiety and Low STEM Anxiety students. Future work should include a larger sample to ensure inclusion of High Anxiety and High Science Anxiety students in connectivity analyses. Second, some participants may have experienced elevated anxiety during the MRI scanning session, potentially confounding results. As this work focuses on the effects of anxiety on functional connectivity, future studies may include MRI-related anxiety as a predictor variable (Ahlander et al., 2016). Third, significantly increased generalized anxiety was observed among High Math Anxiety compared to Low STEM Anxiety students. While the current study found significant STEM anxiety-related differences in DAN, VAN, and DMN connectivity after controlling for generalized anxiety, it is unknown to what extent STEM and generalized anxiety may interact to disrupt functional brain connectivity. Finally, results from this study may not generalize to other groups of university students such as non-STEM students who may experience and adapt to STEM-related anxiety differently due to reduced exposure to STEM courses in their curriculum. Furthermore, since this study examined undergraduate physics students, results may not generalize to other physics cohorts or STEM content domains.

## Conclusions

This study confirmed attentional control theory (ACT) in the context of STEM-related anxiety and demonstrated that undergraduate physics students with elevated STEM anxiety exhibited reduced task-based connectivity among brain networks that collaborate to maintain and modulate attentional control. Specifically, we observed significant and widespread STEM anxiety-related differences within and between the dorsal attention network (DAN), ventral attention network (VAN), and default mode network (DMN) during physics conceptual reasoning and answer selection (i.e., FCI task Phase III), but no differences during problem initiation or question presentation. These results suggest the importance of sustained cognition and DMN-related processing associated with STEM-related anxiety during the FCI task. Further, we found no significant DMN-related differences during the PK task, which measures physics-based content knowledge. Significant group differences during the PK task were limited to the DAN, including DAN-VAN and within-DAN connectivity. These results highlight the different cognitive processes that are required to complete the PK task compared to the FCI task. Finally, we observed no significant differences in FCI or PK task performance between High Math Anxiety and Low STEM Anxiety students. However, it is unclear if greater anxiety would lead to greater variability in task performance or if sustained anxiety experiences can lead to long-term performance differences. Future work is needed to explore the potential effects of STEM-related anxiety across STEM courses or learning experiences to determine if STEM exposure attenuates or exacerbates the task-based connectivity differences observed in the present study. Enhanced insight into the complex relations between STEM-related anxiety and STEM performance will likely yield more effective interventions to improve STEM retention and graduation rates.

## Acknowledgments

Data collection for this project was funded by NSF REAL DRL-1420627 (ARL, EB, SMP, JEB). Contributions from co-authors were provided with support from NSF 1631325 (ARL, MCR, TS), NIH R01-DA041353 (ARL, MTS, MCR), NIH U01-DA041156 (ARL, MTS, MCR, KLB). Special thanks to the FIU Instructional & Research Computing Center (IRCC, http://ircc.fiu.edu) for providing the HPC and computing resources that contributed to the research results reported within this paper, as well as to the Department of Psychology of the University of Miami for providing access to their MRI scanner. Additional thanks to Dr. Jeremy Elman for sharing semantic retrieval task stimuli. Lastly, the authors would like to thank the FIU undergraduate students who volunteered, participated, and contributed to this project.

A Github repository was created at https://github.com/donishadsmith/Physics_Learning to archive all data processing and data analysis scripts used for this study.

All figures used in this study have been archived at https://figshare.com/projects/Figures/149170.

The parcellation and ROI masks that were used in this study are available via Neurovault at https://neurovault.org/collections/12958.

The OSF project page for this study is available at https://osf.io/3w8gt/.

## Author Contributions

DDS, ARL, AM, JEB conceived and designed the study. JEB acquired data. DDS, MCR analyzed data. KLB, MCR, TS, JP contributed scripts, pipelines, and figures. DDS, ARL, AM wrote the paper and all authors contributed to the revisions and approved the final version.

