## Supplemental Information for "Task-based attentional and default mode connectivity associated with STEM anxiety profiles among university physics students"

This document includes:

- Supplemental Methods
- Supplemental Results
- Supplemental Table 1
- Supplemental Figure 1
- Supplemental References

### SUPPLEMENTAL METHODS

#### Neuroimaging Preprocessing

Preprocessing of functional magnetic resonance imaging (fMRI) data was performed using `fMRIPrep` 1.5.0, a BIDS-App that automatically provides a dynamic and adaptable workflow depending on the input data; thus, ensuring high-quality preprocessing with minimal manual intervention (Esteban et al., 2019).

##### Anatomical Data Preprocessing

T1-weighted images were corrected for intensity non-uniformity (INU) with `N4FieldCorrection` (Tustison et al., 2010), distributed with `ANTs` 2.2.0 (Avants et al., 2008, RRID:SCR_004757). The T1w-reference was then skull-stripped with a `Nipype` implementation of the `antsBrainExtraction.sh` workflow (from `ANTs`), using `OASIS30ANTs` as target template. Brain tissue segmentation of the cerebrospinal fluid (CSF), white-matter (WM) and gray-matter (GM) was performed on the brain-extracted T1w using `fast` (`FSL` 5.0.9, RRIDLSCR_002823, Zhang et al., 2001). A T1w-reference map was computed after registration of 2 T1w images (after INU-correction) using `mri_robust_template` (`FreeSurfer` 6.0.1, Reuter et al., 2010). Brain surfaces were reconstructed using `recon-all` (`FreeSurfer` 6.0.1, RRID:SCR_001847, Dale et al., 1999), and the brain mask estimated previously was refined with a custom variation of the method to reconcile `ANTs`-derived and `FreeSurfer`-derived segmentations of the cortical gray-matter of `Mindboggle` (RRID:SCR_002438, Klein et al., 2017). Volume-based spatial normalization to one standard space (MNI152NLin2009cAsym) was performed through nonlinear registration with `antsRegistration` (`ANTs` 2.2.0), using brain-extracted version of both T1w reference and the T1w template. The following template was selected for spatial normalization: ICBM 152 Nonlinear Asymmetrical template version 2009c {Fonov et al. (2009), RRID:SCR_008796: TemplateFlow ID: MNI152NLin2009cAsym].

##### Functional Data Preprocessing

Each participant's dataset contained 1-3 runs of task-based functional magnetic resonance imaging (fMRI) data. For each BOLD run found per subject (across all tasks and sessions), the following preprocessing was performed. First, a reference volume and its skull-stripped version were generated using a custom methodology of `fMRIPrep`. A deformation field to correct for susceptibility distortions was estimated based on *`fMRIPrep`*' s *fieldmap-less* approach. The deformation field is that resulting from co-registering the BOLD reference to the same-subject T1w-reference with its intensity inverted (Huntenburg, 2014; Wang et al., 2017). Registration is performed with antsRegistration (`ANTs` 2.2.0), and the process regularized by constraining deformation to be nonzero only along the phase-encoding direction, and modulated with an average fieldmap template (Treiber et al., 2016). Based on the estimated susceptibility distortion, an unwarped BOLD reference was calculated for a more accurate co-registration with the anatomical reference. The BOLD reference was then co-registered to the T1w reference using `bbregister` (`FreeSurfer`) which implements boundary-based registration (Greve & Fischl, 2009). Co-registration was configured with six degrees of freedom. Head-motion parameters with respect to the BOLD reference (transformation matrices, and six corresponding rotation and translation parameters) are estimated before any spatiotemporal filtering using `mcflirt` (`FSL` 5.0.9, Jenkinson et al., 2002). The BOLD time-series, were resampled to surfaces on the following spaces: *fsaverage5*. The BOLD time-series (including slice-timing correction when applied) were resampled onto their original, native space by applying a single, composite transform to correct for head-motion and susceptibility distortions. These resampled BOLD time-series will be referred to as *preprocessed BOLD in original space*, or just *preprocessed BOLD*. The BOLD time-series were resampled into standard space, generating a *preprocessed BOLD run in ['MNI152NLin2009cAsym'] space*. First, a reference volume and its skull-stripped version were generated using a custom methodology of *fMRIPrep*. Several confounding time-series were calculated based on the *preprocessed BOLD*: framewise displacement (FD), DVARS and three region-wise global signals. FD and DVARS are calculated for each functional run, both using their implementations in *Nipype* (following the definitions by Power et al. 2014). The three global signals are extracted within the CSF, the WM, and the whole-brain masks. Additionally, a set of physiological regressors were extracted to allow for component-based noise correction (*CompCor*, Behzadi et al. 2007). Principal components are estimated after high-pass filtering the *preprocessed BOLD* time-series (using a discrete cosine filter with 128s cut-off) for the two *CompCor* variants: temporal (`tCompCor`) and anatomical (`aCompCor`). `tCompCor` components are then calculated from the top 5% variable voxels within a mask covering the subcortical regions. This subcortical mask is obtained by heavily eroding the brain mask, which ensures it does not include cortical GM regions. For `aCompCor`, components are calculated within the intersection of the aforementioned mask and the union of CSF and WM masks calculated in T1w space, after their projection to the native space of each functional run (using the inverse BOLD-to-T1w transformation). Components are also calculated separately within the WM and CSF masks. For each CompCor decomposition, the *k* components with the largest singular values are retained, such that the retained components' time series are sufficient to explain 50 percent of variance across the nuisance mask (CSF, WM, combined, or temporal). The remaining components are dropped from consideration. The head-motion estimates calculated in the correction step were also placed within the corresponding confounds file. The confound time series derived from head motion estimates and global signals were expanded with the inclusion of temporal derivatives and quadratic terms for each (Satterthwaite et al., 2013). Frames that exceeded a threshold of 0.5 mm FD or 1.5 standardised DVARS were annotated as motion outliers. All resamplings can be performed with *a single interpolation step* by composing all the pertinent transformations (i.e. head-motion transform matrices, susceptibility distortion correction when available, and co-registrations to anatomical and output spaces). Gridded (volumetric) resamplings were performed using `antsApplyTransforms` (`ANTs`), configured with Lanczos interpolation to minimize the smoothing effects of other kernels (Lanczos, 1964). Non-gridded (surface) resamplings were performed using `mri_vol2surf` (`FreesSurfer`).

### SUPPLEMENTAL RESULTS

#### Latent Profile Analysis

Latent Profile Analysis (LPA) was conducted to identify groups of students with similar STEM-related anxiety characteristics. The LPA identified four profiles, including High STEM Anxiety (n=8, 6.5% of the sample), Low STEM Anxiety (n=73, 59.3% of the sample), High Math Anxiety (n=27, 21.9% of the sample), and High Science Anxiety (n=5, 4.1% of the sample). Given low sample sizes for the High STEM Anxiety and High Science Anxiety groups, these profiles were excluded from subsequent analysis. Thus, further analysis only focused on examining differences between the High Math Anxiety and Low STEM Anxiety groups. **Supplementary Table 1** depicts the demographic information for the High Math Anxiety and Low Stem Anxiety groups.

*Table S1*. Demographic Information of the High Math Anxiety and Low STEM Anxiety Groups.

| | Number | Percentage |
|---|---|---|
| **High Math Anxiety** | | |
| **Gender** | | |
| Male | 12 | 44 |
| Female | 15 | 56 |
| **Ethnicity** | | |
| Hispanic | 20 | 74 |
| Non-Hispanic | 7 | 26 |
| **Household Income** | | |
| < $15,000 | 5 | 19 |
| $15,000 - $34,999 | 5 | 19 |
| $35,000 - $49,999 | 3 | 11 |
| $50,000 - $74,999 | 4 | 15 |
| $75,000 - $99,999 | 6 | 22 |
| >$100,000 | 4 | 15 |
| **Years Enrolled** | | |
| Freshman | 2 | 8 |
| Sophomore | 9 | 36 |
| Junior | 10 | 40 |
| Senior | 4 | 16 |
| | **Mean (Std. Dev.)** | **Range** |
| **Age** | 20.0 (1.4) | 18-24 |
| **GPA** | 3.2 (0.4) | 2.3-3.9 |
| **Low STEM Anxiety** | | |
| **Gender** | | |
| Male | 49 | 67 |
| Female | 24 | 33 |
| **Ethnicity** | | |
| Hispanic | 50 | 68 |
| Non-Hispanic | 23 | 32 |
| **Household Income** | | |
| < $15,000 | 18 | 25 |
| $15,000 - $34,999 | 13 | 18 |
| $35,000 - $49,999 | 11 | 15 |
| $50,000 - $74,999 | 12 | 16 |
| $75,000 - $99,999 | 8 | 11 |
| >$100,000 | 11 | 15 |
| **Years Enrolled** | | |
| Freshman | 6 | 9 |
| Sophomore | 38 | 58 |
| Junior | 16 | 25 |
| Senior | 5 | 8 |
| | **Mean (Std. Dev.)** | **Range** |
| **Age** | 19.5 (1.5) | 18-26 |
| **GPA** | 3.3 (0.6) | 0.0-4.0 |

**Supplementary Figure 1** shows the distribution of the normalized science anxiety and math anxiety scores for the High Math Anxiety and Low STEM Anxiety groups.

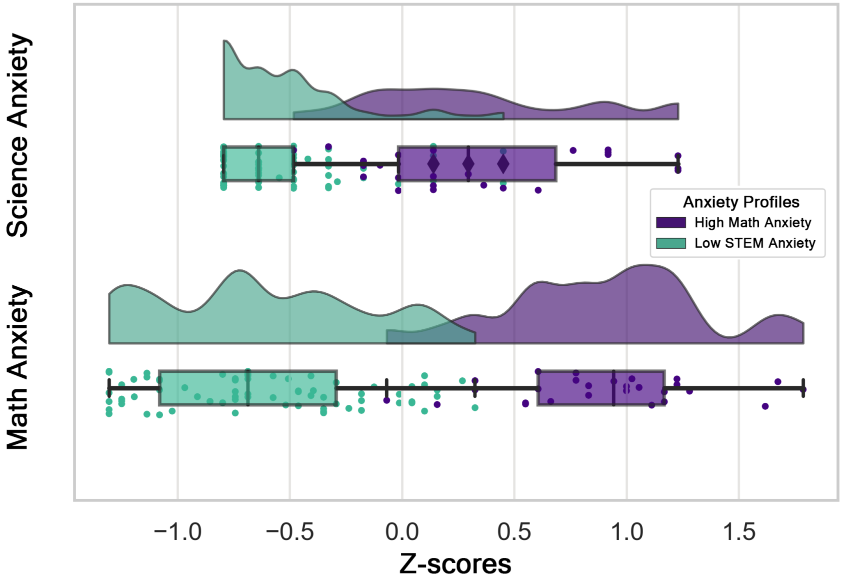


*Figure S1.* **Science and Math Anxiety Scores for High Math Anxiety and Low STEM Anxiety Groups.** Distribution of the normalized science and math anxiety scores in the High Math Anxiety (purple) and Low STEM Anxiety (green) groups.
